# RNA sequence analysis of somatic mutations in aging and Parkinson’s Disease

**DOI:** 10.1101/2025.03.26.645360

**Authors:** Shixiang Sun, Daisy Sproviero, César Payán-Gómez, Jan H.J. Hoeijmakers, Alexander Y. Maslov, Pier G. Mastroberardino, Jan Vijg

## Abstract

Parkinson’s Disease (PD) is an age-related neurodegenerative disorder that has been associated with increased DNA damage. To test if PD is associated with increased somatic mutations, we analyzed RNA-seq data in whole blood from 5 visits of the Parkinson’s Progression Markers Initiative for clonally amplified somatic variants. Comprehensive analysis of RNA-sequencing data revealed a total of 5,927 somatic variants (2.4 variants per sample on average). Mutation frequencies were significantly elevated in PD subjects as compared to age-matched controls at the time of the last visit. This was confirmed by RNA analysis of substantia nigra. By contrast, the fraction of carriers with clonal hematopoiesis, was significantly reduced in old PD patients as compared to old healthy controls. These results indicate that while the overall mutation rate is higher in PD, specific clonally amplified mutations are protective against PD, as has been found for Alzheimer’s Disease.

## Introduction

Parkinson’s disease (PD) is the second most common neurodegenerative disorder, affecting 2–3% of the population over 65 years old ^1^. While age is the greatest risk factor, PD also has a genetic component with about 4% of cases due to single-gene defects and genetic variation estimated to contribute about 25% to the overall risk of developing this complex disorder ^2^. The etiology of PD is multifactorial with two major pathological hallmarks: the selective loss of dopaminergic neurons of the substantia nigra (SN), and the presence of Lewy bodies containing α-synuclein ^1^. Evidence has been obtained that the molecular basis of PD involves mitochondrial dysfunction leading to an increased production of reactive oxygen species (ROS) causing death of dopaminergic neurons ^3^. It has been proposed that this ROS-induced cell death and dysfunction is mediated by increased oxidative DNA damage, which may be accelerated by partial defects in DNA repair ^4,5^. Moreover, the cellular stress, mediated by α-synuclein has been shown to elicit DNA damage ^6^.

An increasingly studied molecular end point of the interaction between DNA damage and repair in neurodegenerative and other complex genetic diseases is somatic mutation. Many if not most genetic diseases have a somatic mutational counterpart due to clonally amplified somatic mutations arising during early development ^7,8^. In PD, sporadic cases could be caused by somatic mutations in the same genes affected in familial cases ^9^. Indeed, in Alzheimer’s disease (AD), somatic mutations in autosomal dominant genes, such as amyloid precursor protein (APP) and presenilin 1 (PS-1), have been found in brain samples as causes of sporadic forms of the disease, albeit only in few cases ^10^. In these studies, increased somatic mutations were also found in blood ^10^. In PD, evidence for clonally amplified somatic mutations has been recently obtained for different regions of the brain as well as in blood, affecting genes enriched in synaptic and neuronal functions ^11^. Such clonally amplified somatic mutations could play critical functional roles if present in expressed genes ^12^. Here we tested for clonally amplified somatic mutations in transcribed sequences as a possible risk factor for PD through a mutation calling pipeline in RNA-seq data from blood and SN.

## Results

### PD-related accumulation of somatic mutations in blood

To test for clonally amplified somatic mutations in PD subjects at the mRNA level, we used RNA-seq data from blood of 179 controls and 385 PD patients available through the Parkinson’s Progression Markers Initiative (PPMI; **Fig. 1A**) ^13^. In the PPMI cohort, the ages of subjects at visits were between 31 and 85.2 years (**Table S1**). To evaluate the progression of PD, blood samples for RNA-seq were collected at five time points, ranging from the first visit to follow-up visits at 6, 12, 24 and 36 months (visits were named BL, V02, V04, V06 and V08; **Fig. 1B**), bringing the total number of blood samples to 2,443 (**Table S1**). Of note, the numbers of subjects in PPMI were not the same from visit to visit because not all subjects were sequenced at each visit (**Fig. 1B**). All PD patients enrolled in the PPMI study had been diagnosed with PD with demonstrated dopaminergic deficits. Of the PPMI PD patients, 41 and 9 were carriers of the pathogenic germline variants in glucosylceramidase beta (GBA) or leucine rich repeat kinase 2 (LRRK2) genes, respectively. There were no significant differences in age (P = 0.468) or sex (P = 0.925) between PD patients and healthy controls.

**Fig. 1.**
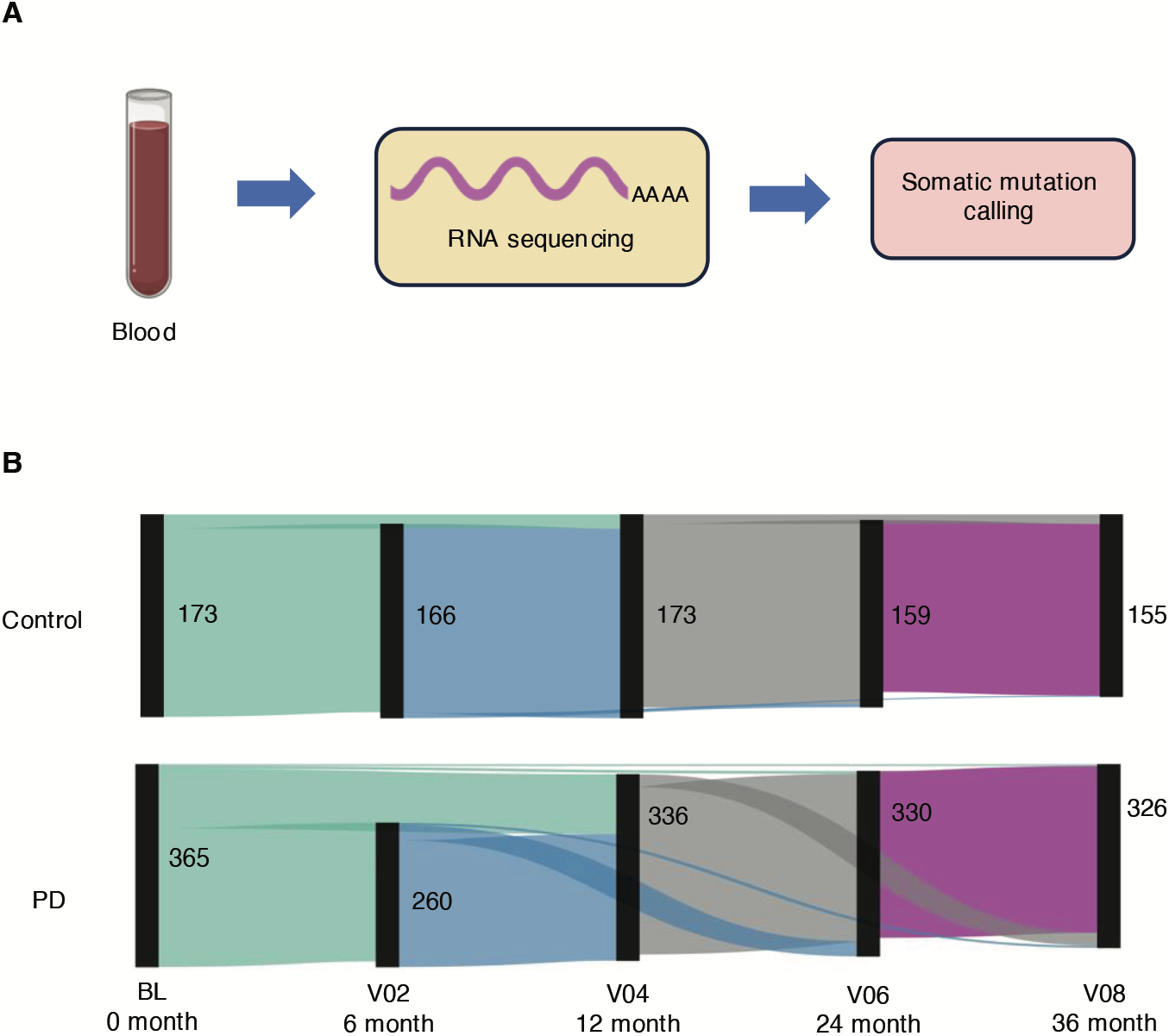
Study overview. (A) Workflow of sequencing and data processing. Sequencing data was obtained from 4 publications and PPMI. This figure is created with the help of BioRender. (B) Sankey diagram of sequenced subjects in five visits at 0 months (baseline), 6 months, 12 months, 24 months, and 36 months. The numbers next to the bars indicate the total number of samples sequenced in that visit, while horizontal bands show the samples sequenced in two connected visits.

To characterize the mutation profiles in whole blood RNA-seq, we applied the RNA-MuTect-WMN pipeline, which can detect single nucleotide variants (SNVs) in protein-coding genes and their promoters at high precision and high sensitivity (Methods; **Table S2**) ^14^. A variant was considered a true mutation when it was shown in at least 4 reads. After analyzing all 2,443 samples a total of 5,927 somatic variants (2.4 variants per sample on average) were found.

In both PD patients and control subjects, variants in blood were found to overlap between visits within the same subject, with on average 89.6% of mutations for a subject (control or PD) called at least twice across all time points (**Fig. S1A**). The percentage of recalled mutations was found to decline from 1 to 3 years after the baseline visit at 75.3%, 70.6% and 63.5% in controls, as well as 70.4%, 65.1% and 63.3% in PD patients (V04, V06 and V08, respectively), suggesting a gradual loss of clonally amplified mutations over a 2-year period. Overlap of mutations in the same subjects across time points as well as a gradual loss of mutations over time are expected since mutational variants are likely to persist in blood cells over a period of time but will eventually disappear due to slow loss of entire mutant cell lineages.

In contrast to mutations called within each subject, 95.9% and 96.1% of mutated genomic positions were unique among the different healthy controls and different PD patients, respectively (**Fig. S1B**), compared with 97.3% unique mutated positions in healthy subjects from the GTEx study ^15^. These results indicate that clonally amplified mutations called from RNA are vastly different between subjects.

Next, we compared mutation burden in PD patients to age-matched healthy controls. The results for 5 visits showed that the average SNV frequency per individual in PD patients was elevated compared with controls (1.08, 1.01, 1.08, 1.04 and 1.16-fold; **Fig. 2**), but this was only statistically significant for the last visit, i.e., V08 (P = 0.048, **Fig. 2E**).

**Fig. 2.**
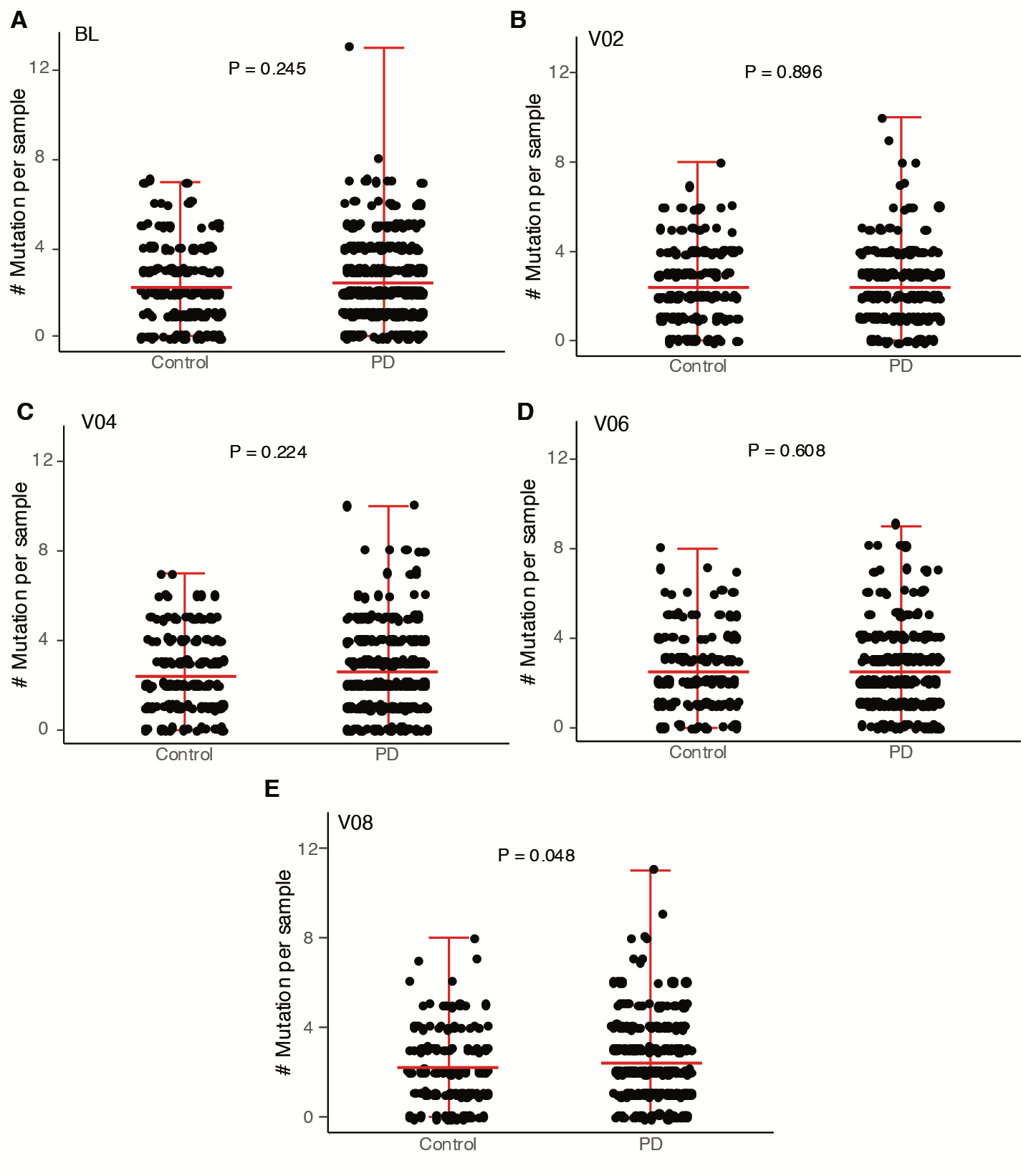
Numbers of somatic mutations in controls and PD cases. The number of mutations in (A) BL, (B) V02, (C) V04, (D) V06, and (E) V08. Each data point indicates the observed numbers in each subject. The red line in the middle indicates the mean number of mutations for each age group, while the top and bottom red lines are the minimal and maximal range of mutations.

We then tested if the elevated somatic mutation frequencies in blood could be confirmed in affected brain tissue of PD patients. For this purpose, we used the same variant calling strategy to study somatic mutations at the mRNA level in SN of PD patients and controls using four public RNA-seq datasets ^16-19^. We collected RNA-seq data from 24 healthy controls and 32 PD subjects. The ages of subjects at death were between 61 and 95 years, with no significant differences in age (P = 0.095). A gender difference was also absent (P = 0.789). The post-mortem delays for all SN samples were within 24 hours. We identified 83 mutations in the 58 SN samples. As in blood, mutations were found mostly unique between different subjects (94.9%), supporting a high diversity of mutations among the subjects. The results indicate a significantly elevated mutation burden in SN of PD subjects as compared to SN from healthy controls (1.95-fold, P =7.06e-3; **Fig. S2**). The more pronounced increase of somatic mutations in SN as compared to blood coincides with other PD-related changes, most notably changes in levels of Synuclein Alpha, which are more dramatic in postmortem brain samples than antemortem blood ^20^. Overall, this result confirms an elevated burden of somatic mutations in PD patients.

### Variant allele fractions (VAFs) in different cohorts among five visits

We then checked if the variant allele fraction differed between PD patients and control subjects. Here VAF difference is calculated with the average VAFs between PD and control subjects. For each visit, the average VAFs were not significantly increased in PD patients compared with in controls (**Fig. S3A**). However, albeit small, the VAF difference between PD and controls became larger from the first to the fifth visit (R-squared = 0.958, P = 3.57e-3; **Fig. S3B**). This suggests that compared with controls, in PD the blood cells carrying these somatic mutations are further clonally amplified than in controls, implying a possible selective advantage. In addition, we observed a trend towards a higher average VAF in PD patients than in controls in SN, albeit this was also not significant (P = 0.768; **Fig. S3A**).

### Associations between somatic mutation levels and germline variants

There are a number of pathogenic germline variants known to increase PD risk ^2^. Here we specifically analyzed germline variants in LRRK2 and GBA within blood samples. Surprisingly, with the limited number of carriers in this PPMI cohort, the presence of the two germline variants showed reduced somatic mutation frequency in PD patients at all visits (0.90, 0.58, 0.80, 0.77 and 0.80-fold; **Fig. S4**), which was significant for V02 and V06 (**Fig. 3**). The presence of LRRK2 was found associated with lower mutation frequencies than GBA (0.60, 0.83, 0.94, 0.70 and 0.66-fold; **Fig. S4**), albeit this was not statistically significant. LRRK2 is highly expressed in immune cells ^21^ and this PD variant has been found associated with neuronal cell death ^22^. These results suggest that PD-related pathogenic germline variants are unrelated to increased DNA damage as a risk factor.

**Fig. 3.**
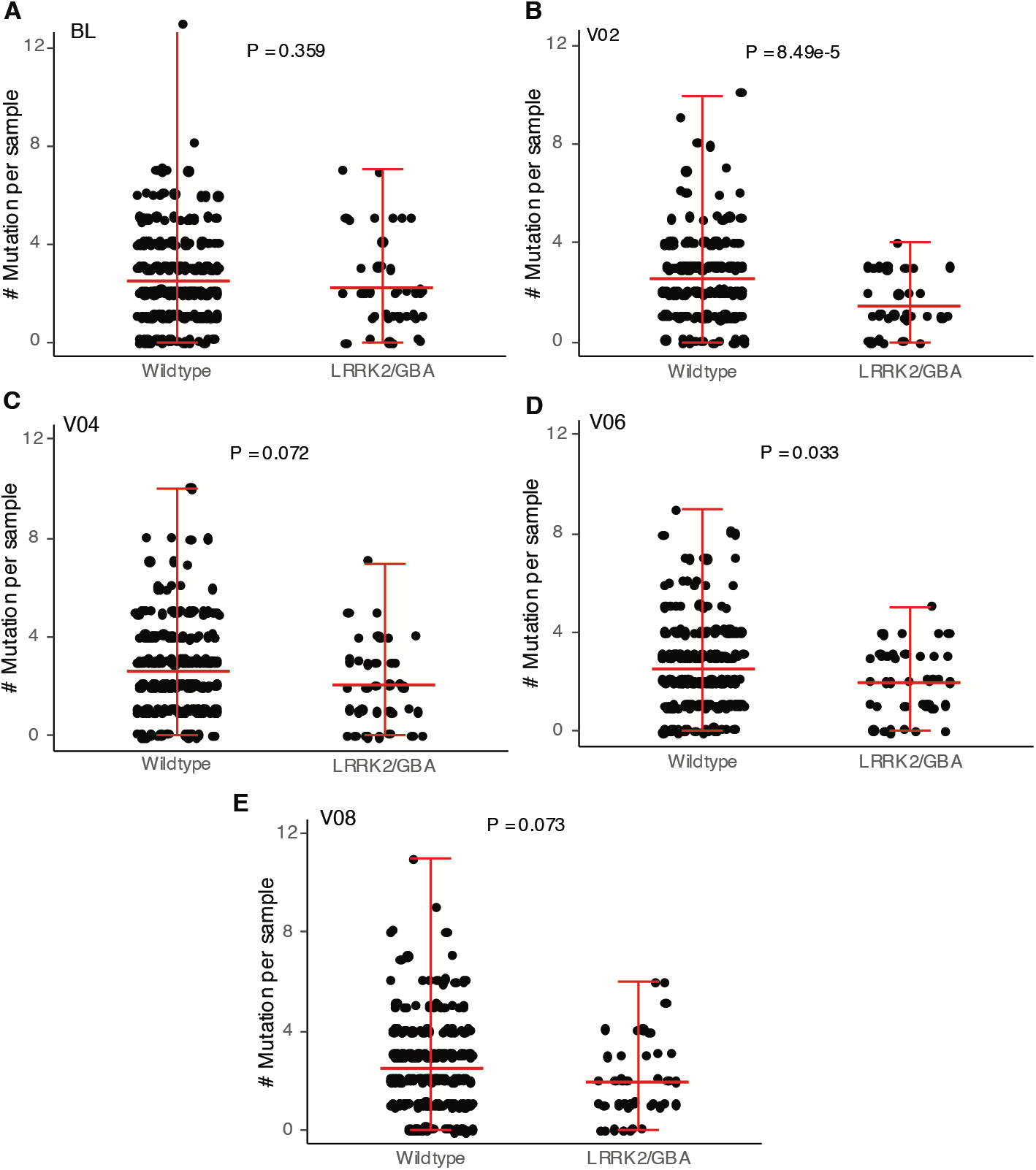
Numbers of somatic mutations in PD cases in relation to the germline variants. The number of mutations in (A) BL, (B) V02, (C) V04, (D) V06, and (E) V08. Each data point indicates the observed numbers in each subject. The red line in the middle indicates the mean number of mutations for wildtype and germline variants carrier group, while the top and bottom red lines are the minimal and maximal range of mutations.

### Reduced frequency of clonal hematopoiesis in PD patients

Since pathogenic germline variants were not associated with an increase in SNV frequency in PD patients, we next investigated whether pathogenic somatic mutations exhibit a distinct pattern in PD as the SNV frequency increased. Recently, the frequency of Clonal Hematopoiesis of Indeterminate Potential (CHIP) carriers in AD patients was reported to be lower than in age-matched control subjects ^23^, suggesting a protective effect. Here, we hypothesized that the presence of PD can protect the patients against accumulation of pathogenic somatic mutations. To test this, we examined known, cancer-associated CHIP variants in 73 genes in the whole exome sequencing data of our cohort (Methods). We identified CHIP events in 25 subjects (**Table S3, Fig. 4A**).

**Fig. 4.**
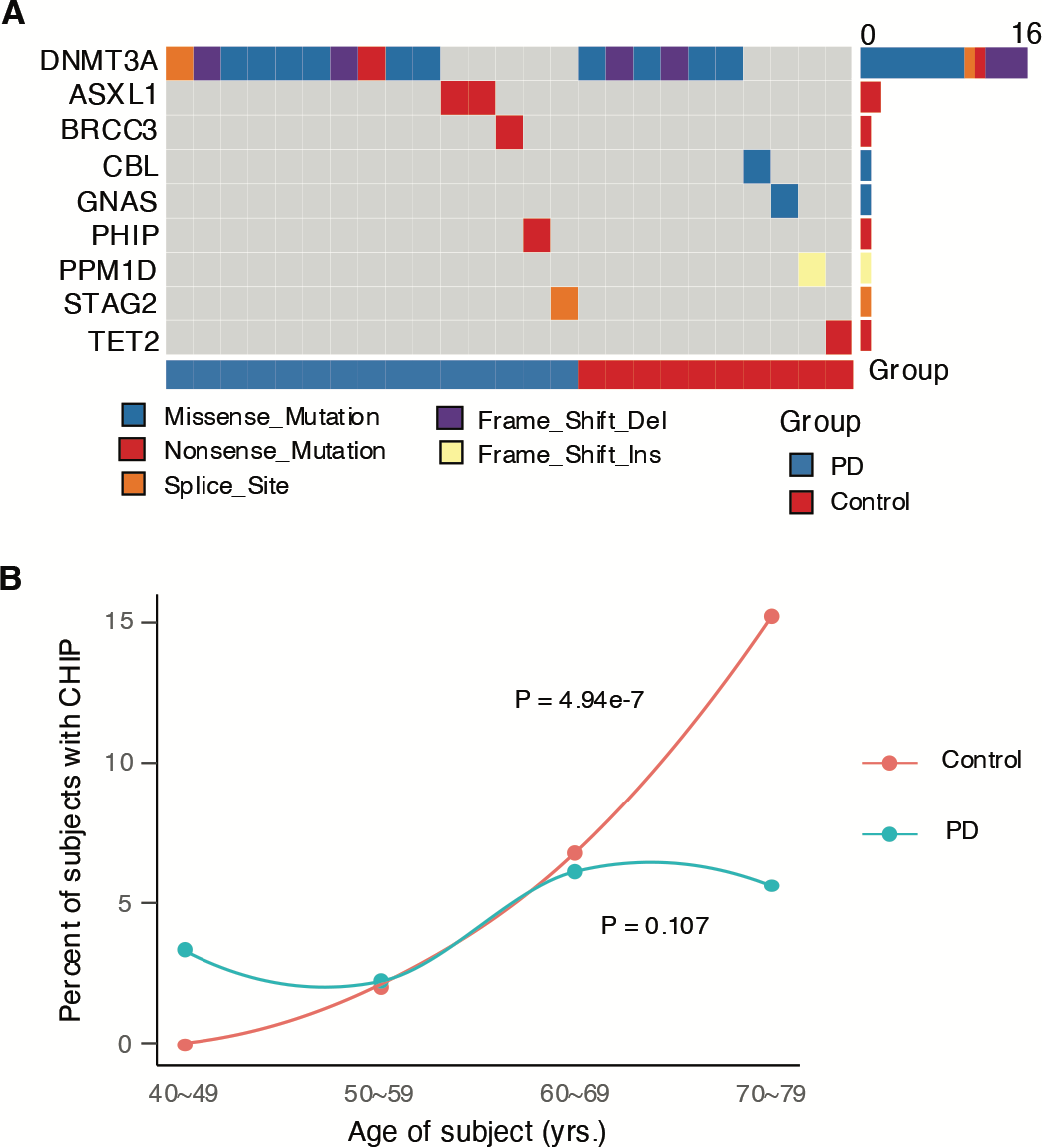
CHIP analysis of WES data in healthy controls and PD subjects. (A) Co-mutation plot with mutations across the 25 subjects from healthy controls and PD patients. (B) Distribution of percentages of CHIP carriers across different age bins.

We then examined if the percentage of CHIP carriers was associated with age or PD status.

As only 5% of the subjects were CHIP carriers, we summarized the results by age bins (**Fig. 4B**). The results indicated that the percentages of CHIP carriers were significantly increased with age in controls (P = 4.94e-7) but not in PD samples (P = 0.107), confirming the well-documented age-related increase of CHIP in blood ^24^. Further, we found that the relationship between age and the percentages of CHIP carriers were significantly different between the healthy controls and PD patients (P = 1.06e-3). As age increases, the likelihood of acquiring CHIP variants was lower in PD patients compared to healthy controls, leading to a lower percentage of CHIP carriers among the oldest age group of PD patients (Ratio = 0.37, P = 0.020). This observation is in keeping with the reported lower CHIP frequencies in AD ^23^. Hence, these results support the notion that PD is protective against specific clonally amplified mutations, which is in keeping with the observed negative association of PD with blood disease ^25^.

### Differences in somatic mutation signatures between PD subjects and healthy controls

To explore the possible source of the somatic mutations detected in the cohort in relation to PD, we analyzed the mutation signatures. We grouped samples according to disease (PD or control) and tissue (SN or blood) into 12 sub-groups and fitted known mutational signatures (Methods). Of the eight identified signatures, we found two signatures SBS7b and SBS31 were uniquely present in PD patients (**Fig. 5, Table S4**).

**Fig. 5.**
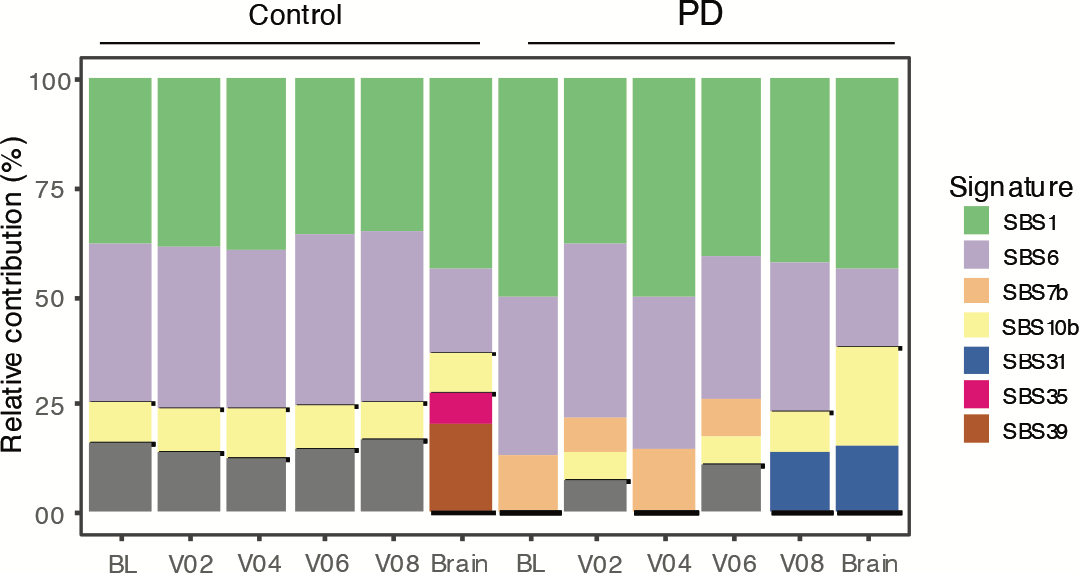
Somatic mutational signatures in the cohort. Figures indicate the contributions of signatures of COSMIC to all SNVs in 12 sub-groups of the surveyed cohorts. Bars marked with brain are SN samples.

SBS7b was only present in first four visits of blood in PD patients, and SBS31 was only found in SN and last visit of blood from PD patients (**Fig. 5**). SBS7b was found in CD4+ T lymphocytes ^26^, while SBS31 was observed in pediatric brain tumors ^27^. These two signatures were both featured GC to AT transitions and associated with transcription-coupled nucleotide excision repair on DNA damage ^28^. They were similar to each other (cosine similarity: 0.74), and the major two mutational subtypes C[C>T]C and C[C>T]T were shared between the signatures (**Fig. S6**). Additionally, SBS35 was only found in SN control samples and present similar etiology as SBS31, but GC to TA and AT to TA transversions were the dominations in SBS35, which were not in SBS31.

Signature SBS1 was found significantly increased in the PD patient group compared with the healthy control group in SN and blood together (control vs. PD on average: 38.6% vs. 44.2%, P = 0.049; **Fig. 5**). SBS1 is known as age-related signatures ^28^. This result indicates that aging signature SBS1 are also accumulated in PD patients. This is not surprising because PD is an age-related disease ^29^. Interestingly, signature SBS1 has been associated with oxidative damage ^30^, which has been causally implicated in both aging and PD.

## Discussion

Using a highly sensitive and precise pipeline^14^ we called clonally amplified somatic mutations in blood RNA-seq data from the ongoing observational PD cohort study PPMI, consisting of 5 visits over a time period of 3 years. This allowed us to characterize the landscape of somatic mutations in regions of the protein-coding genes from whole blood of PD patients as compared to age-matched healthy controls. Our results showed that SNV frequencies were increased in blood from PD patients, albeit this was significant only at the last visit. These results were confirmed by analyzing RNA-seq data from SN samples, which also revealed significantly higher SNV frequencies in PD patients than in healthy controls.

The most straightforward way of interpreting our results is based on the hypothesis that increased oxidative stress, for example, caused by mitochondrial dysfunction, leads to increased DNA damage, which causes the elevation of mutation frequency observed in PD patients. This would assume that any such mitochondrial dysfunction is systemic and affects not only the brain but also the blood. Such a possibility would be in keeping with the observed increase of somatic mutation frequency in both blood and SN of PD patients ^11^.

Alternatively, the increased somatic mutation frequency could be explained by a minor deficiency in one or more DNA repair pathways. Major defects in DNA repair have been found associated with accelerated aging in both mice and humans ^31^. However, not much is known about minor DNA repair defects, not affecting any of the core DNA repair processes in a major way. Nevertheless, it is not at all unlikely that reported DNA repair defects in neurodegenerative diseases are based on such minor defects. Elevated levels of oxidative stress due to mitochondrial dysfunction and minor DNA repair defects are of course not mutually exclusive.

The above interpretation would be in keeping with the observed mutational spectra. The oxidative damage associated GC to AT transitions or mutational signatures featuring this mutation type were found to be unique to PD (SBS7b and SBS31) or increased in PD (SBS1) compared with healthy controls. Of note, SBS7b was found in the first four visits and SBS31 in the last visit, suggesting changes over time. SBS31 was also found in SN samples.

It is generally assumed that the causes of PD involve a combination of genetic and environmental factors. In this respect, the possibility should be considered that the reported correlation between PD and DNA repair defects ^32^ points towards one of multiple risk factors rather than a major causal relationship. When PD etiology would increase oxidative stress, DNA damage and mutations would also increase. This effect would be more severe if DNA repair activities in a subject are on the low side. This could well explain the long-standing implication of DNA damage and DNA repair defects in PD etiology. This possibility is in keeping with our observation that the pathogenic germline mutations in LRRK2 and GBA do not seem to increase somatic mutation burden in PD patients, suggesting different, unrelated risk factors.

Somewhat unexpectedly, the observed higher somatic mutation frequencies in PD were not reflected in a higher CHIP frequency. Instead, CHIP frequency was lower among PD patients. Hence, like AD *(25)*, also PD is negatively associated with the accumulation of the pathogenic mutations in blood. This is in contrast to previously observed associations of CHIP levels with risk of age-related, chronic disease, most notably cardiovascular disease and overall mortality ^33^, and suggest that like AD also PD protects against CHIP or CHIP protects against PD. In the AD study *(25)*, the evidence suggests that CHIP-mutated clones infiltrate the brain and prevent the onset of AD. Since we had no access to PD brain samples from PPMI cohort, we cannot rule out that a similar process may be associated with the onset of PD. Beyond CHIP, PD has previously been found negatively associated with multiple cancers, including leukemia ^25^, which is also consistent with the inverse associations found between AD and cancers ^34^. Thus, it is possible that the lower cancer risk is due to PD patients developing less CHIP clones.

In conclusion, our results indicate a more complex relationship between DNA damage metabolism and PD than previously assumed. While a systemic increase in somatic mutations indicate adverse DNA damage effects, possibly through increased oxidative damage, those clonally amplified somatic mutations that have been found associated with blood cancer are reduced in PD, as they are in AD.

## Methods

### Data source

Information of surveyed subjects in blood was obtained from PPMI (file names: meta_data.11192021.csv and Age_at_visit.csv; https://ida.loni.usc.edu/). The gender information missing from the PPMI files were taken from ^35^. Raw RNA and whole exome sequencing data in blood were downloaded from the PPMI’s server. Specifically, before sequencing, whole-transcriptome RNA was extracted from blood cells after rRNA and globin reduction. Raw RNA sequencing data in brain SN samples were downloaded from SRA (**Table S1**), and sample information were collected from the corresponding publications.

### Alignment of the sequencing data

Raw RNA sequencing data were trimmed for adapter and low-quality nucleotides with Trim Galore (version 0.6.7). The minimal length of reads for alignment was set at 30 bp. Clean RNA-seq data were aligned with STAR (version 2.6.1d) ^36^, with pre-defined parameters ^15^. The build of the reference genome was GRCh37.

Raw whole exome sequencing data were trimmed for adapter and low-quality nucleotides with Trim Galore (version 0.6.7). The minimal length of reads for alignment was required to be at least 30 bp. Alignments were performed according to the best practices of the Genome Analysis Toolkit (GATK version: 4.2.5.0) ^37^. In general, trimmed reads were aligned with BWA (mem mode; version 0.7.17) ^38^. Removal of PCR duplications and base recalibration were performed in the alignment files using GATK.

### Calling somatic mutations

Somatic mutations were called from the RNA-seq data following the pipeline RNA-MuTect-WMN ^14^. In general, mutations were first identified using Mutect (version 1.1.6) ^39^ with the ‘-U ALLOW_N_CIGAR_READS’ flag. Following the pipeline, only mutations found in exons, splicing regions, and promoters (5kb upstream of the transcription start sites) of a protein-coding gene were maintained for further analysis (listed in gaf_20111020.broad_wex_1.1). Somatic candidates were identified using RNA-MuTect-WMN (RNA-Mutect-WMN-test.py), employing the random forest classifier for separating germline and somatic variants. According to RNA-MuTect-WMN, the somatic candidate mutations were then filtered using the filtering steps in RNA_MUTECT ^15^. The default parameters and reference files are collected as ^40^. First, mutation candidates were annotated using GATK Oncotator (version 1.8.0.0; data source: oncotator_v1_ds_Jan262014). Reads covering candidates were realigned with Hisat2 (version 2.0.3) ^41^ and mutations were recalled and confirmed with Mutect. Rest filtering steps were performed using scripts in RNA_MUTECT with default settings (run_FilterRNAMutations.sh and run_FilterRNAMutationsBasedOnDuplicateReads.sh). Here, the panel of normal files were collected from GTEx ^42^ and TCGA ^43^. Only variants present in at least 4 reads were denoted as true mutations.

### Examining CHIP variants

To examine the association between CHIP and PD, we excluded the subjects with known PD pathogenic germline variants or unknown phenotypes. Subjects younger than 40 years old or older than 80 years old at consent are excluded, as too few samples are in these two bins. Mutect2 (version 4.2.5.0) ^44^ was applied to call known variants in known CHIP associated genes obtained from ^45^. Candidates were kept with adjusted pre-applied filters ^46^: if they passed Mutect2 default filtering, and had a read depth ≥20x, the number of reads supporting variant alleles ≥7 in single nucleotide variants and ≥10 in other mutation types, the variant allele fraction (VAF) ≥2%, gnomAD allele frequency <0.001, and at least one read in both forward and reverse direction supporting the reference and variant alleles.

### Identifying mutation signatures

To refit the known mutation signatures, we used non-negative least-squares. To avoid overfitting and removal signatures with little effects, we applied strict refitting using function ‘fit_to_signatures_strict’ in R package MutationalPatterns ^47^. The input data were 12 subgroups grouped by their PD status, visits and tissues. Known mutations were collected from COSMIC ^48^ if they are present in originally publication ^28^. Eight mutational signatures are fitted for the 12 subgroups of the samples The cosine similarity was calculated to compare different signatures.

### Statistical analyses

To compare the distribution of age and sex between PD and healthy controls, two-tailed student’s t-test and Fisher’s exact test were used, respectively. A negative binomial generalized linear model was applied to estimate the effects of PD status and carrying pathogenic germline mutations, with the number of observed mutations as the response and age bin at visit and sex as fixed effects, offset by the surveyed genome coverage. The significances were assessed using the likelihood ratio test. A two-way ANOVA was applied to test the differences in average VAFs between PD and controls. For CHIP analysis, a generalized linear regression model was applied to examine the effect of age or PD status, including intersection of PD status on age bin increase, with likelihood ratio test to assess the statistical significance. Exact binomial test was used to compare the percentage of CHIP carriers in oldest groups of healthy control and PD subjects. The differences of relative contributions on mutation signatures between PD patients and controls were tested with paired t-test.

## Acknowledgments

We thank all members of the Jan Vijg’s laboratory for helpful discussions related to this project. We thank the staffs in the PPMI for assisting the data downloading and medication logs access.

Data used in the preparation of this article were obtained on [2023-02-23] from the Parkinson’s Progression Markers Initiative (PPMI) database (https://www.ppmi-info.org/access-data-specimens/download-data), RRID:SCR_006431. For up-to-date information on the study, visit http://www.ppmi-info.org.

PPMI – a public-private partnership – is funded by the Michael J. Fox Foundation for Parkinson’s Research and funding partners, including 4D Pharma, Abbvie, AcureX, Allergan, Amathus Therapeutics, Aligning Science Across Parkinson’s, AskBio, Avid Radiopharmaceuticals, BIAL, BioArctic, Biogen, Biohaven, BioLegend, BlueRock Therapeutics, Bristol-Myers Squibb, Calico Labs, Capsida Biotherapeutics, Celgene, Cerevel Therapeutics, Coave Therapeutics, DaCapo Brainscience, Denali, Edmond J. Safra Foundation, Eli Lilly, Gain Therapeutics, GE HealthCare, Genentech, GSK, Golub Capital, Handl Therapeutics, Insitro, Jazz Pharmaceuticals, Johnson & Johnson Innovative Medicine, Lundbeck, Merck, Meso Scale Discovery, Mission Therapeutics, Neurocrine Biosciences, Neuron23, Neuropore, Pfizer, Piramal, Prevail Therapeutics, Roche, Sanofi, Servier, Sun Pharma Advanced Research Company, Takeda, Teva, UCB, Vanqua Bio, Verily, Voyager Therapeutics, the Weston Family Foundation and Yumanity Therapeutics.

## Funding

This study was supported by The Michael J. Fox Foundation (J.V., P.G.M.); NIH grants P01 AG017242, P01 AG047200, P30 AG038072 and U01 ES029519, the Glenn Foundation for Medical Research (J.V.).

## Author contributions

J.V. and P.G.M. conceived and supervised the study. S.S. analyzed the data. S.S., D.S., C.P.G., J.H.H, A.Y.M, P.G.M., and J.V. wrote the manuscript.

## Competing interests

A.Y.M and J.V. are the co-founders of SingulOmics, Corp. Other authors declare no competing interests.

## Data and materials availability

Raw sequencing data in blood, including RNA-seq data and whole exome sequencing data were obtained from https://www.ppmi-info.org. Raw sequencing data in brain were obtained from SRA under accession numbers SRP255916, SRP219862, SRP382117, and SRP202025. The raw mutation calling pipeline is available in https://github.com/yizhak-lab-ccg/RNA_MUTECt_WMN, with the docker files from https://hub.docker.com/r/noamrud/rna_mutect_wmn_test and https://hub.docker.com/r/kyizhak/rna_mutect. Additional data and script related to this paper may be requested from the authors.

## Notes

### Summary of Updates

We updated data acknowledgment and the names of the PPMI industry partners in the acknowledgements section

