## Supplementary figure for "RNA sequence analysis of somatic mutations in aging and Parkinson’s Disease"

### Supplementary Figures

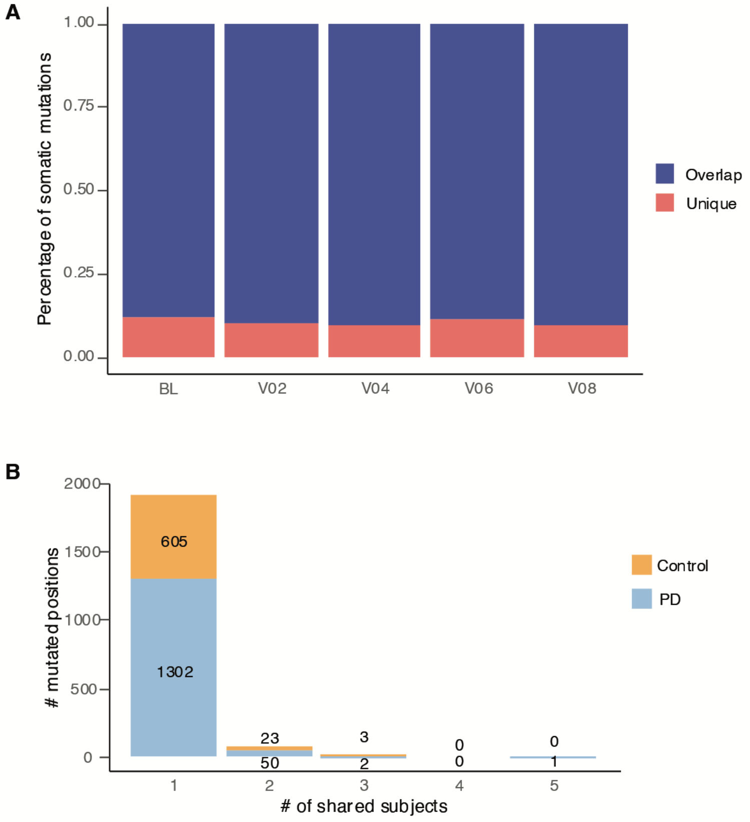


**Fig. S1. Overlapping mutations.** (A) Percentage of mutations overlapping in each visit of blood samples. (B) number of mutations shared by different subjects in the PPMI cohort.

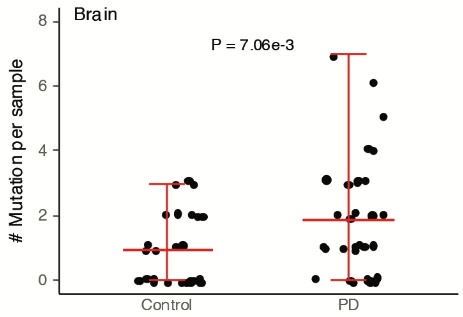


**Fig. S2. Numbers of somatic mutations in SN samples.** Number of mutations are plotted. Each data point indicates the observed numbers in each subject. The red line in the middle indicates the mean number of mutations for each age group, while the top and bottom red lines are the minimal and maximal range of mutations.

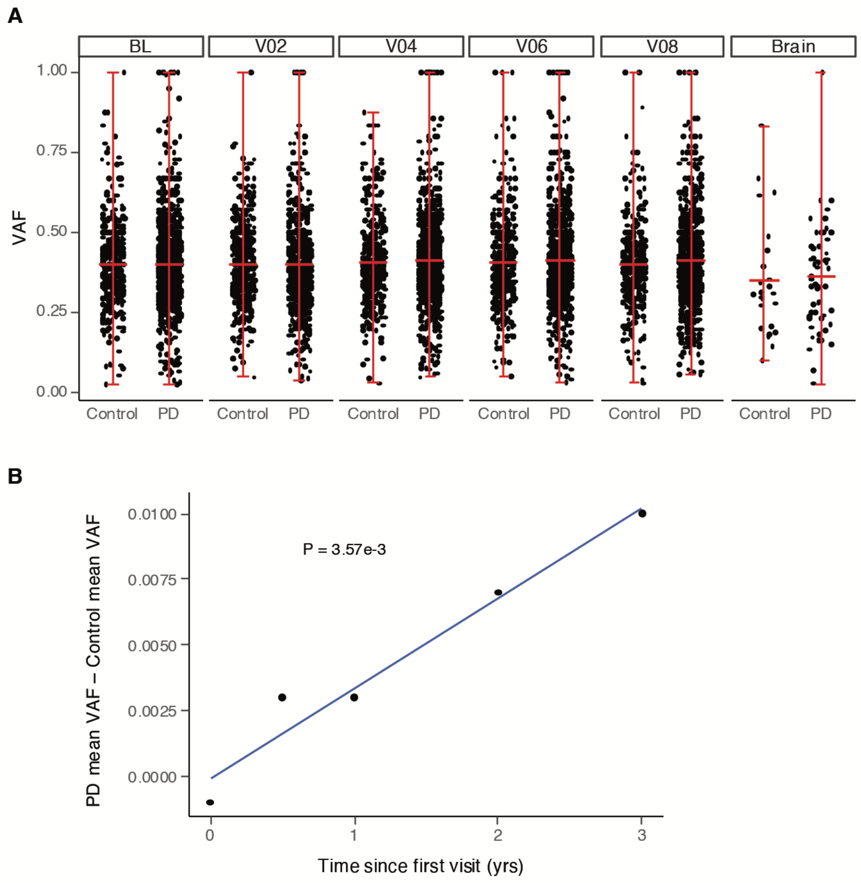


**Fig. S3. Variant allele frequencies (VAFs) of somatic mutations.** (A) The VAFs are plotted for SN and blood (each visit), separated by PD status. Each data point indicates the observed numbers in each subject. The red line in the middle indicates the mean number of mutations for each age group, while the top and bottom red lines are the minimal and maximal range of mutations. (B) delta VAF between PD patients and controls across the 5 visits.

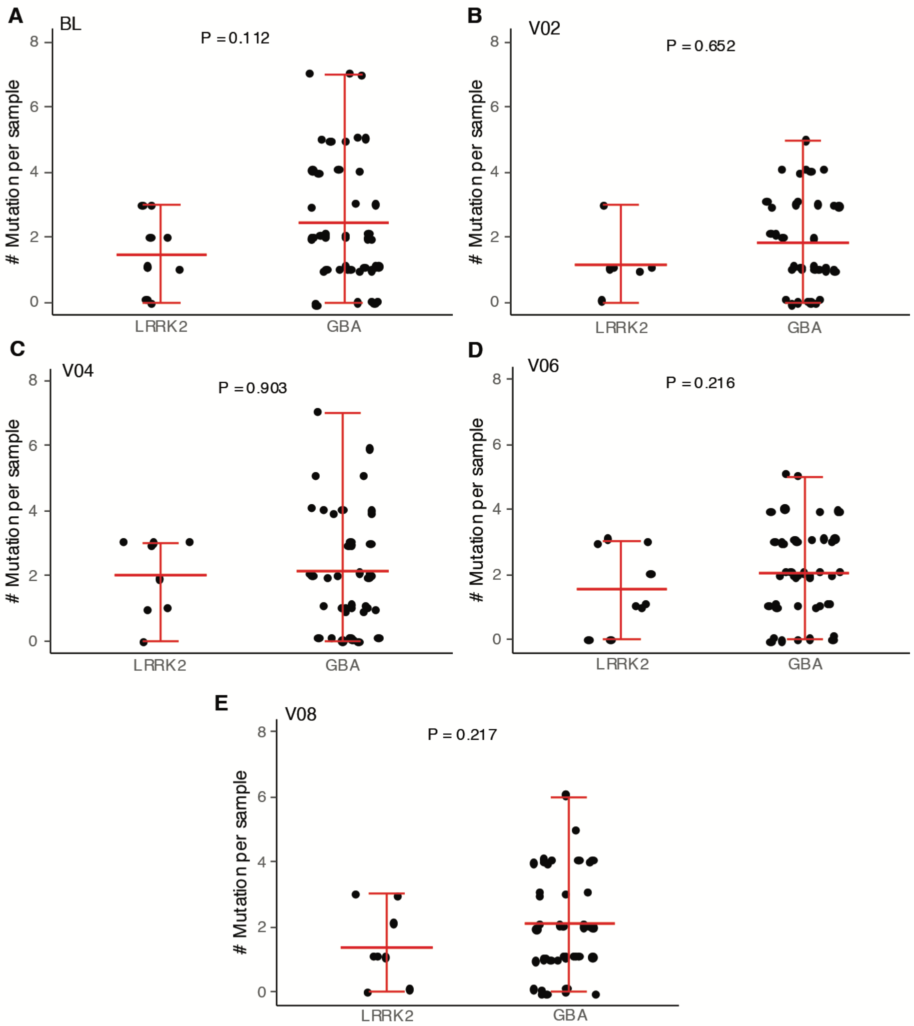


**Fig. S4. Numbers of somatic mutations in different pathogenic germline carriers of PD cases.** The number of mutations in (A) BL, (B) V02, (C) V04, (D) V06, and (E) V08. Each data point indicates the observed numbers in each subject. The red line in the middle indicates the mean number of mutations for LRRK2 and GBA carrier groups, respectively, while the top and bottom red lines are the minimal and maximal range of mutations.

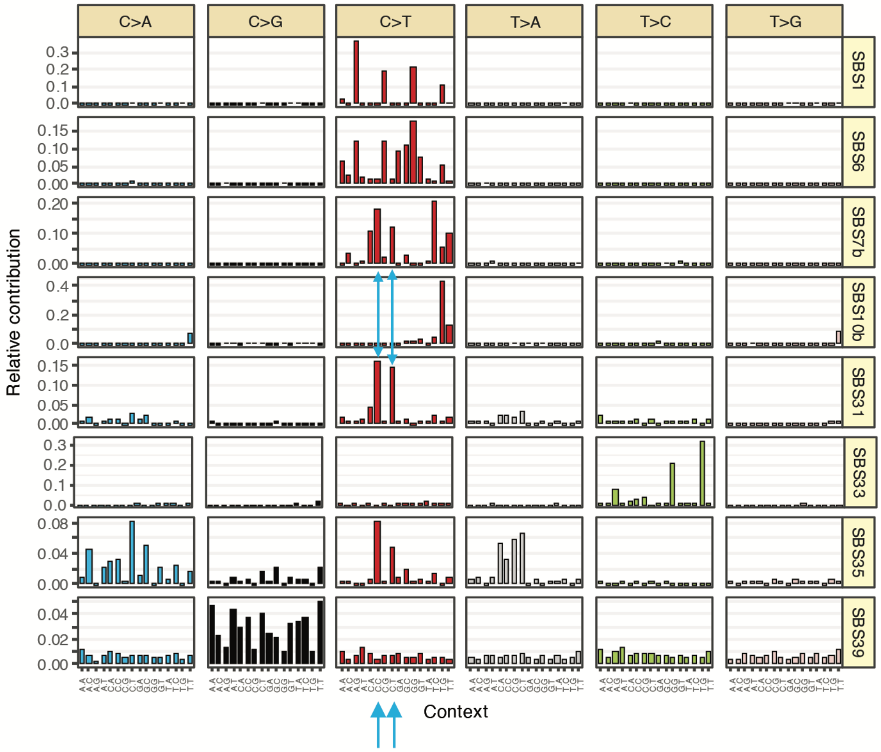


**Fig. S5 Mutational spectrum of fitted COSMIC signatures.** Two overlapping mutation types between SBS7b and SBS31 are marked by blue arrows.
